# Design and performance of a long-read sequencing panel for pharmacogenomics

**DOI:** 10.1101/2022.10.25.513646

**Authors:** Maaike van der Lee, Loes Busscher, Roberta Menafra, Qinglian Zhai, Redmar R. van den Berg, Sarah B Kingan, Nina Gonzaludo, Ting Hon, Ting Han, Leonardo Arbiza, Ibrahim Numanagić, Susan L. Kloet, Jesse J. Swen

## Abstract

Pharmacogenomics (PGx)-guided drug treatment is one of the cornerstones of personalized medicine. However, the genes involved in drug response are highly complex and known to carry many (rare) variants. Current technologies (short-read sequencing and SNP panels) are limited in their ability to resolve these genes and characterize all variants. Moreover, these technologies cannot always phase variants to their allele of origin. Recent advance in long-read sequencing technologies have shown promise in resolving these problems. Here we present a long-read sequencing panel-based approach for PGx using PacBio HiFi sequencing.

A capture based approach was developed using a custom panel of clinically-relevant pharmacogenes including up- and downstream regions. A total of 27 samples were sequenced and panel accuracy was determined using benchmarking variant calls for 3 Genome in a Bottle samples and GeT-RM star(*)-allele calls for 21 samples..

The coverage was uniform for all samples with an average of 94% of bases covered at >30×. When compared to benchmarking results, accuracy was high with an average F1 score of 0.89 for INDELs and 0.98 for SNPs. Phasing was good with an average of 68% the target region phased (compared to ~20% for short-reads) and an average phased haploblock size of 6.6kbp. Using Aldy 4, we compared our variant calls to GeT-RM data for 8 genes (*CYP2B6, CYP2C19, CYP2C9, CYP2D6, CYP3A4, CYP3A5, SLCO1B1, TPMT*), and observed highly accurate star(*)-allele calling with 98.2% concordance (165/168 calls), with only one discordance in *CYP2C9* leading to a different predicted phenotype.

We have shown that our long-read panel-based approach results in high accuracy and target phasing for SNVs as well as for clinical star(*)-alleles.

## 1 Introduction

Pharmacogenomics (PGx) is one of the main pillars of personalized medicine, promising better treatment outcomes by adjusting drug therapy based on the individual’s genetic make-up (1). While PGx is increasingly applied in clinical practice, the complexities of the genes involved (pharmacogenes) limit our ability to accurately predict drug response (2, 3). From twin studies it is known that 60-90% of drug metabolism by cytochrome P450 enzymes is heritable (4–7). However, currently applied tests usually focus on a limited set of relatively common genetic variants and can only explain 30-50% of the observed variability (4–7).

Three of the major challenges are the complexity of pharmacogenes, the presence of low frequency and rare variants, and haplotype phasing. It is known that the majority of pharmacogenes are at least partially complex, where complex is defined as containing duplicated regions or high sequence similarity to other genes or to pseudogenes (3, 8). For example, the *CYP3A* genes (*CYP3A4, CYP3A5* and *CYP3A7*) share approximately 85% of their sequence (9). Depending on the genotyping technology used, this makes it difficult to fully distinguish between these genes. Secondly, pharmacogenes are known to be extremely polymorphic, with many rare variants (minor allele frequency (MAF) <1%). While some low frequency and rare variants are expected to have a major impact on protein function, they are often neglected in routine PGx diagnostics (10, 11). Currently used tests often focus on common and pre-defined variants and are unable to identify low frequency and rare variants. Finally, haplotype phasing might play a role in our inability to explain all genetic variability in drug response. Without phasing, it is unclear which variants are located on the same allele and which are on the opposite allele. This becomes of clinical importance if a patient carries two deleterious variants. If these variants are on the same allele there may be one non-functional and one functional allele, resulting in a decreased overall protein function. However, if the variants are on opposing alleles there may be no functional allele present, resulting in a decrease or absence of protein function. Due to the high number of variants in the pharmacogenes, phasing is expected to be especially important in this context.

Currently, next-generation sequencing (NGS) using short reads (~150-200bp) and single nucleotide variant (SNV)-based approaches are most commonly used in PGx research and clinical practice. These approaches are able to accurately call variants and, in the case of NGS, identify novel and rare variants. However, calling (novel) variants in regions that are difficult to resolve due to sequence homology or repetitive sections remains a challenge. Long-read sequencing applications hold promise to resolve complex regions into large and phased haploblocks (12). With PacBio HiFi sequencing, more than 6.5kbp can be sequenced in one read when using a capture-based approach. To increase the accuracy of this sequence, each sequenced molecule undergoes several passes, yielding a highly accurate consensus sequence (12).

However, for PGx, the application of long-read sequencing has been limited to either a single gene or single patient approach using existing whole genome sequencing (WGS) data (13–16). We believe that a PacBio HiFi long-read sequencing panel-based approach can be valuable for PGx research. In this paper we describe the design and first performance of such a HiFi sequencing-based PGx panel.

## 2. Methods

All work described in this paper is subject to the LUMC Good Research Practice & Integrity guidelines and Ethical requirements.

### 2.1 Sample selection

Three genome-in-a-bottle (GIAB) samples consisting of an Ashkenazim child-parent trio were used for the protocol development and accuracy assessment at the Leiden Genome Technology Center (LGTC). Upon finalization of the protocol, a further 24 samples were sequenced at the PacBio facilities. Pharmacogenetics results from the Genetic Testing Reference Materials Coordination Program (GeT-RM) are available for 21 of these samples.

### 2.2 Panel selections

The genes in this panel were selected based on their presence in the Dutch Pharmacogenomics Working Group (DPWG) guidelines, supplemented with genes which were of research interest. For each gene, a region of approximately 10kbp upstream and 10kbp downstream was included. Target regions were sent to Twist Bioscience (South San Francisco, CA, USA) for capture probe design.

### 2.3 gDNA Library Preparation

The concentration of the high molecular weight (HMW) genomic DNA (gDNA) used in this study was assessed using the Qubit Fluorometer and the Qubit dsDNA Broad Range Assay kit (Invitrogen, Carlsbad, CA). The integrity of the HMW gDNA was verified on the Femto Pulse system using the Genomic DNA 165 kb Kit (Agilent Technologies, CA, USA. Three microgram of HMW gDNA was sheared to an average size distribution of ~8 kb using the Covaris g-TUBE (Covaris, MA, U.S.A.) in an Eppendorf 5810 R centrifuge at 5,000 rpm for a total of 2 minutes. Before continuing the protocol, AMPure XP beads (Beckman-Coulter, IN, USA) were washed according to the PacBio protocol “Guidelines for using AMPure XP Beads for HiFi sequencing.” These washed beads were used to clean and concentrate sheared DNA with a 0.5X v/v ratio followed by a concentration measurement using the Qubit Fluorometer and the Qubit dsDNA High Sensitivity Assay kit (Invitrogen, Carlsbad, CA). The fragment size distribution was verified on the Femto Pulse System using the Genomic DNA 165 kb Kit.

The 24 Coriell samples prepared at PacBio follow the same workflow with the following modifications: sheared genomic DNA was concentrated using a 0.6X v/v ratio of AMPure PB beads and libraries were sized using a Bioanalyzer 2100 instrument with the DNA 12000 Kit (Agilent Technologies, CA, USA).

### 2.4 Target Capture

500 ng of sheared and purified gDNA was used as input for end repair, A-tailing and adapter ligation using the Twist Library Preparation Mechanical Fragmentation Kit 1 (Twist Bioscience, CA, USA) according to the manufacturer’s protocol “Twist Mechanical Fragmentation and Twist Universal Adapter System (doc-001087 rev 2.0, April 2021).” The Twist adapters were replaced with single-stranded linear barcoded adapters (10 μM stock, Integrated DNA Technologies (IDT), IA, USA) which were annealed according to the PacBio protocol “Multiplex Genomic DNA Target Capture Using IDT xGen Lockdown Probes (Version 04, July 2018).”

Adapter-ligated sample was purified using washed AMPure XP beads with a 0.8X v/v ratio followed by size selection using 3.7X diluted washed AMPure XP beads (35% w/v) and elution into 51 μl of 10 mM Tris-HCl, pH 8.5. Next, pre-capture amplification was performed using the Takara LA Taq HotStart kit (TaKaRa Bio USA, Inc.) in two, 100 μL reaction volumes containing 50–100 ng of adapter-ligated DNA with a final concentration of 0.5 μM PacBio universal primer (/5Phos/gcagtcgaacatgtagctgactcaggtcac (IDT, IA, USA)), 0.1 mM of each dNTP, 1x LA PCR buffer, and 0.03 units/μL Takara LA HotStart Taq. The PCR parameters were 2 minutes at 95°C followed by 6 cycles of 20 seconds at 95°C, 15 seconds at 64°C, and 10 minutes at 68°C with a final extension of 5 minutes at 68°C. After amplification, the two reaction volumes were combined and samples were purified with 0.5X v/v washed AMPure XP beads and eluted in 20 μl of 10 mM Tris-HCl, pH 8.5. The DNA concentration and size distribution of the amplified sample was assessed using the Qubit Fluorometer with the Qubit dsDNA High Sensitivity Assay kit and the Femto Pulse System using the Genomic DNA 165 kb Kit, respectively. After quality control, samples were pooled in equimolar amounts with a maximum of eight samples in one pool.

After pooling, the samples were captured using the Twist Hybridization and Wash Kit (Twist Bioscience, CA, USA) and a Twist probe custom panel (Twist Bioscience, CA, USA) according to the Twist Target Enrichment Standard Hybridization v1 Protocol (Revision 2.0, May 2021) with the following changes: the PacBio universal primer was used instead of the Twist universal blockers and Dynabeads M-270 Streptavidin (Invitrogen, CA, USA) were used instead of Twist streptavidin binding beads. The captured pools were amplified using the Takara HotStart kit in two, 100 μl reaction volumes containing 50 μl captured sample pool and a final concentration of 0.5 μM PacBio universal primer, 0.2 mM of each dNTP, 1x LA PCR buffer, and 0.03 U/μL Takara LA HotStart Taq. The PCR parameters were 2 minutes at 95°C followed by 13 cycles of 20 seconds at 95°C, 15 seconds at 64°C and 10 minutes at 68°C with a final extension of 5 minutes at 68°C. After amplification, the samples were purified with 0.5X v/v washed AMPure XP Beads and eluted in 40 μl of 10 mM Tris-HCl, pH 8.5. The DNA concentration and size of the amplified, captured samples were assessed using the Qubit Fluorometer and the Qubit dsDNA High Sensitivity Assay kit, and the Femto Pulse System and the Genomic DNA 165 kb Kit, respectively.

The 24 Coriell samples prepared at PacBio follow the same workflow with the following modifications: 200 ng of sheared gDNA was used as input for adapter ligation and size selection after adapter ligation was performed using 3.1X v/v of 35% AMPure PB beads. Both pre- and post-capture amplification were performed in one, 100 μL reaction using 50 μL of eluted sample, 5 units of Takara LA Taq (TaKaRa Bio USA, Inc.) and 1 μL of 100 μM PacBio universal primer. The PCR parameters for pre-capture amplification were 2 min at 95°C followed by 10 cycles of 20 sec at 95°C, 15 sec at 62°C, and 10 min at 68°C with a final extension of 5 min at 68°C. The post-capture PCR parameters were 2 min at 95°C followed by 18 cycles of 20 sec at 95°C, 15 sec at 62°C, and 10 min at 68°C with a final extension of 5 min at 68°C. After amplification, samples were purified using 0.5X v/v AMPure PB beads and eluted in 51 μL of EB.

### 2.5 SMRTbell Library Construction and Sequencing

For the GIAB-LUMC samples, sequencing libraries were prepared using 500 ng of pooled samples following the “Procedure & Checklist – Preparing SMRTbell Libraries using PacBio Barcoded Overhang Adapters for Multiplexing Amplicons (version 02, April 2020)” with the SMRTbell Express Template Prep Kit 2.0. The SMRTbell library was sequenced on a Sequel II platform using Sequencing Primer V4, Sequel II Sequencing Kit 2.0 and Sequel II Binding Kit 2.0 on an 8M SMRT cell with a 30 hour movie time. The SMRTbell library was loaded at an on-plate concentration of 80 pM. For the Coriell-PacBio samples, sequencing libraries were prepared using the SMRTbell Prep Kit 3.0 with Sequel II Binding Kit 3.2 and Sequel II Sequencing Kit 2.0.

### 2.6 Processing

A publicly available pipeline developed at the LUMC was used to process the HiFi sequencing data (https://github.com/biowdl/PacBio-subreads-processing) and to call variants (https://github.com/LUMC/PacBio-variantcalling)(17). LIMA v2.2.0 (https://lima.how) was used to demultiplex the circular consensus (CCS) reads and duplicate reads were marked using pbmarkdup v1.0.2 (https://github.com/PacificBiosciences/pbmarkdup). Reads were aligned to GRCh38 using pbmm2 v1.3.0 following which Deepvariant v1.0.0 (18) was used for variant calling and WhatsHap v1.0 for phasing (19). Statistics and quality control (QC) parameters were reported with a custom MultiQC plug-in (20). Parameters reported include: percentage of the target region phased, percentage of duplicate reads, percentage of target bases with < 30x coverage and the total number of variants identified. Results from MultiQC and VCF files were processed using Rstudio (v 1.3.1056) and Python 3.9.12. Benchmarking was performed for the three GIAB samples using Hap.py v3.12.1 (https://github.com/Illumina/hap.py) and v4.2.1 of the NIST benchmarks (21)

For clinical star (*)-allele calling, all samples were analyzed using Aldy4 with the custom setting ‘pacbio-targeted-hifi-twist’, details of which are described by Hari et al (22, 23). Results were compared to the reference calls from GeT-RM, who provide genetic testing reference materials for PGx. Only the major alleles are reported.

## 3. Results

### 3.1 Coverage and accuracy

The samples had an average of 117k HiFi reads each with a mean on-target read length of ~6.5 kb. The percent of target bases covered at least 30x was fairly uniform, averaging 94% per sample with the exception of samples NA18518 (87%) and NA18868 (89%) (Figure 1A). The number of unique molecules was high (mean=85%) and the number of duplicate reads low (mean = 15%) (Figure 1B).

**Figure 1:**
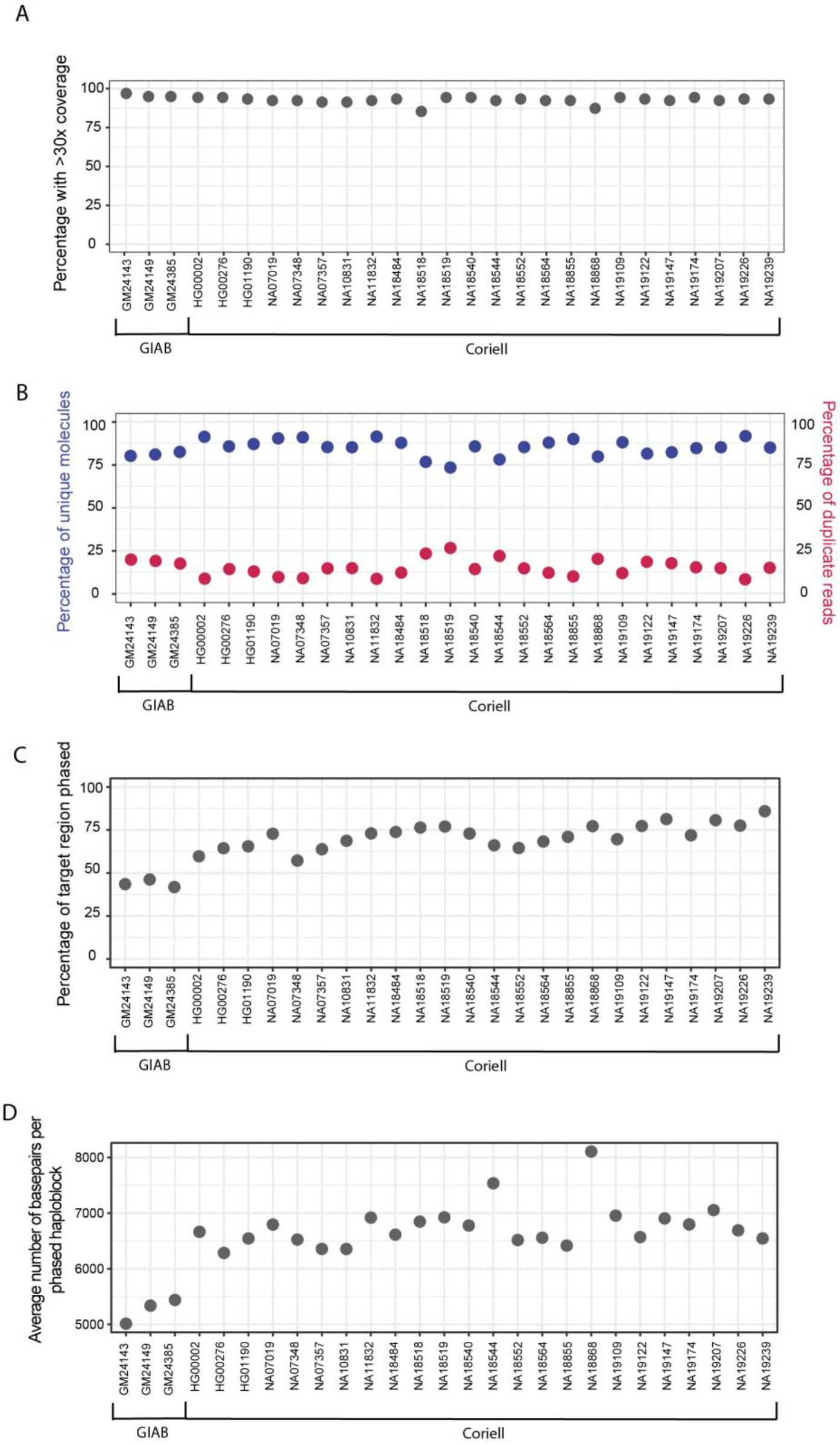
General capture panel performance. (A) On target coverage was high in all samples. (B) The number of duplicated reads was low and the percentage of unique molecules high. (C) An average of 68% of the target region was phased and (D) on average each phased haploblock was 6.6kbp.

Variant benchmarking was performed with the NIST truth sets for the GIAB Ashkenazim trio using hap.py. Overall variant-calling accuracy was high (>86% for recall and precision), with better performance on SNPs (F1 score >0.97) compared to INDELs (F1 score >0.88) (Table 1). Variant density was 4.1 variants per kbp on average, which corresponds to one variant every 244 base pairs. The most variable regions in the panel are *HLA-A* (20.7 variants / kbp) and *CYP2D6* (3.7 variants /kbp) whereas *VKORC1* (1.2 variants / kbp) and *CYP2C19* (1.4 variants / kbp) were relatively well conserved.

**Table 1:** Accuracy compared to benchmarking for Genome in a Bottle trio. INDEL: Insertion or deletion, SNP: Single Nucleotide Polymorphism.

|  | Type | Recall | Precision | F1 score |
| --- | --- | --- | --- | --- |
| GM24149 | INDEL | 0.92 | 0.86 | 0.89 |
|  | SNP | 0.99 | 0.99 | 0.99 |
| GM24385 | INDEL | 0.87 | 0.88 | 0.88 |
|  | SNP | 0.95 | 0.99 | 0.97 |
| GM24143 | INDEL | 0.89 | 0.89 | 0.89 |
|  | SNP | 0.95 | 1.00 | 0.98 |

**Table 2:** Concordance between Aldy 4 and GeT-RM results. The 8 genes reported are: *CYP2B6, CYP2C19, CYP2C9, CYP2D6, CYP3A4, CYP3A5, SLCO1B1, TPMT*

| Sample | Concordance | Remarks |
| --- | --- | --- |
| HG00276 | 100% |  |
| HG01190 | 100% |  |
| NA07019 | 100% |  |
| NA07357 | 100% |  |
| NA10831 | 100% |  |
| NA11832 | 87.5% | CYP2D6*1/*68+*4 was detected as *1/*4 by Aldy |
| NA18484 | 100% |  |
| NA18518 | 100% |  |
| NA18519 | 100% |  |
| NA18544 | 100% |  |
| NA18552 | 100% |  |
| NA18564 | 100% |  |
| NA18855 | 100% |  |
| NA18868 | 100% |  |
| NA19109 | 100% |  |
| NA19122 | 100% |  |
| NA19147 | 100% |  |
| NA19174 | 87.5% | CYP2D6*4/*40 was detected as *4+*4/ *40 by Aldy |
| NA19207 | 100% |  |
| NA19226 | 87.5% | CYP2C9 *1/*8 was detected as *1/*1 by Aldy |
| NA19239 | 100% |  |
| **Total** | **98.2% (165/168)** |  |

### 3.2 Phasing

After phasing reads with WhatsHap, the average on-target phasing was 68% (range 42% - 86%, Figure 1C), compared to an average on-target phasing of ~20% for short-reads on the same target genes (data not shown). Moreover, the average phased haploblock was 6.6kbp long (Figure 1D). When stratifying the phasing per gene a difference was observed, with some genes (e.g. *CYP2B6* and *CYP2C19*) almost completely phased for all samples while for other genes (e.g. *CYP2C9*) only a very small part of the gene is phased into haploblocks (Figure 2A). On average 54% of each gene region could be phased, with the best results observed for *CYP2B6* (84%) and *CYP2C19* (81%). When focusing on only the core variants, thereby ignoring the upstream and downstream regions, phasing remains roughly the same at 51% (Figure 2B). Notably, for some genes, the phasing improved substantially when focusing on the core region. For example, for *TPMT* only 66% of the entire region could be phased, however, on average 89% of the core region was fully phased.

**Figure 2:**
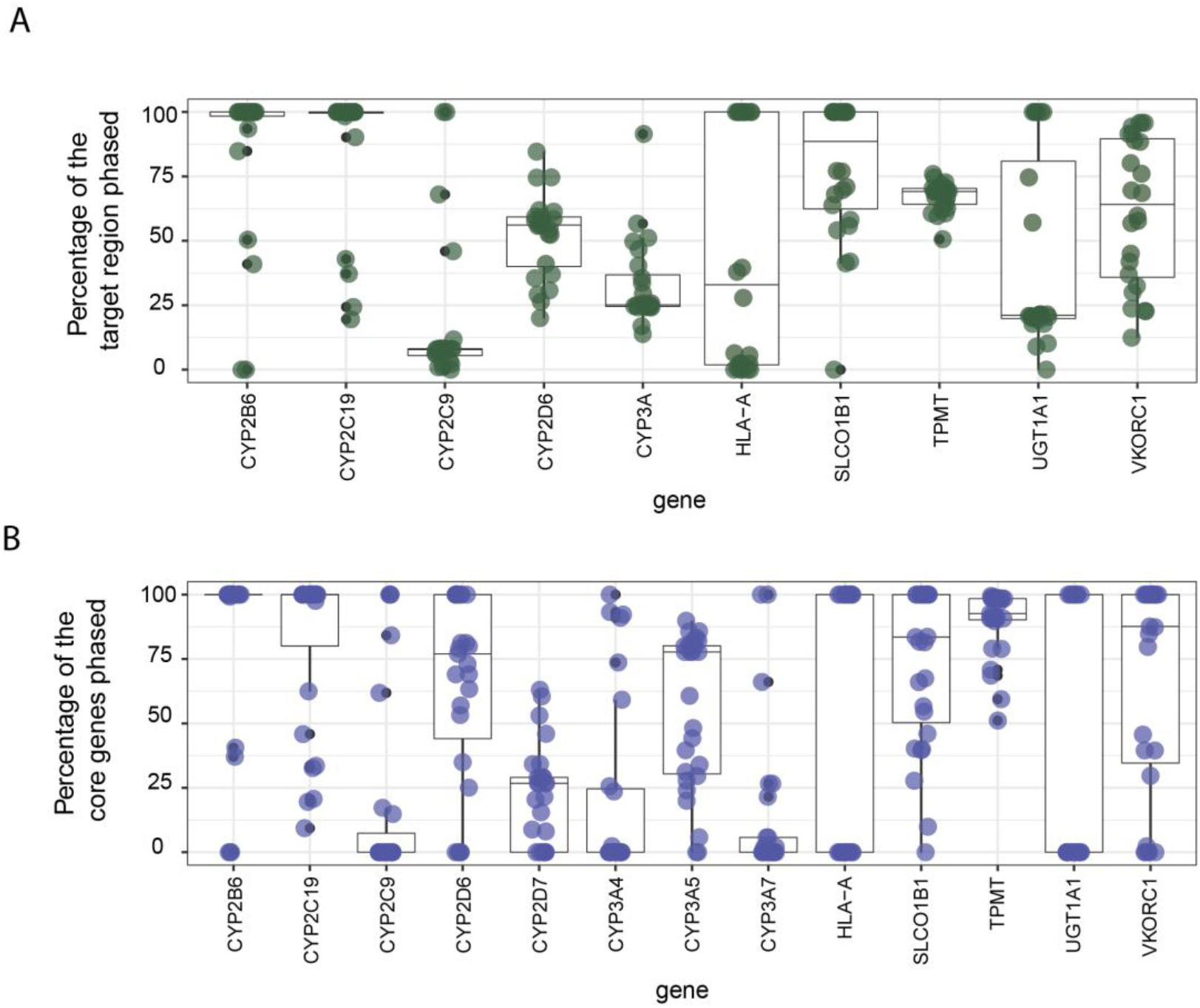
On target phasing of each gene. (A) Phasing performance across the entire target region including up- and downstream regions and (B) phasing performance of the core genes focusing only on the gene body.

For complex regions such as the *CYP3A* locus, consisting of *CYP3A4, CYP3A5* and *CYP3A7*, phasing was poor (30%). When stratifying per gene, the poor phasing remained for *CYP3A4* (21%) and *CYP3A7* (13%) while the phasing of *CYP3A5* (56%) was better resolved (Figure 2B). However, between each of the *CYP3A* genes there was a lack of coverage in the intergenic regions resulting in broken haploblocks and an inability to phase the variants in *CYP3A5* in relation to those in *CYP3A4*.

### 3.3 Performance compared to a clinical standard

To compare to the current clinical standard, 21 samples for which reference data from GeT-RM was available were processed using Aldy 4 to phase the reads and assign the best major (star)*-alleles from the existing star-allele databases. Accuracy was high compared to the GeT-RM curation with 98.2% of the calls (165 out of 168) matching completely based on the major star(*)-alleles. Only one discrepancy (*CYP2C9* * 1/* 1 vs CYP2C9 * 1/*8) would have led to another predicted metabolic phenotype. Inaccuracies were mainly observed for *CYP2D6-CYP2D7* hybrids/conversions (two out of three discrepancies respectively).

## 4. Discussion

Here we present a PacBio HiFi long-read sequencing panel-based approach for pharmacogenomics, which allows high resolution characterization of complex pharmacogenes and phased haplotypes. The developed method results in highly accurate variant calls (>95% recall and precision for SNPs) and relatively high on target phasing (68%). The variant density (average of one variant per 244 basepairs) highlights the importance of long reads for haplotype phasing. Short reads (~150-200bp) will not be able to cover the average distance between two variants and therefore cannot resolve them in one haploblock.

Compared to short-read WGS approaches, the per-gene phasing of our targeted panel was lower (12). However, the use of a panel-based approach costs only a fraction of the price and has a shorter turnaround time compared to WGS. This makes it more accessible for use in clinical practice and in research projects using large cohorts. When compared to short-read panel-based approaches, the presented approach is superior for both accuracy and phasing. Generally, the on-target phasing with short-reads is ~20% while our long-read approach resulted in an average of 68% of bases phased which is substantially higher. Notably, a short-read panel-based approach developed in-house for the same genes only reached F1 scores of 0.94 (SNPs) and 0.85 (INDELs) (data not shown), indicating that the long-read approach is also superior for the variant calling accuracy (F1 scores of 0.98 and 0.89 for SNPs and INDELs respectively).

Nonetheless, not all genes could be fully phased. The main reasons for haploblocks to break are a lack of coverage and a lack of heterozygous variants. With probe optimization it might be possible to improve the phasing for those regions where haploblocks breakage is due to a lack of coverage. The breaking of haploblocks due to homozygosity is however difficult to prevent as this is inherent to each sample’s genetic make-up. Strikingly, for many genes the percentage of the region that could be resolved in one haploblock was larger when focusing on only the core gene region and ignoring the up- and downstream regions (e.g *CYP2C19, SLCO1B1, TPMT*). One potential explanation for this is the higher number of variants in the core genetic region compared to the up- and downstream regions. However, the average variant density is similar in the core regions and the intergenic regions in the panel, making this an unlikely explanation. Another cause could be the spacing between the variants as haploblocks break when encountering large stretches of homozygosity. The panel was designed to not be preferentially targeted towards coding regions but to have even coverage across all bases in the target region.

Haplotypes, as assigned by Aldy 4, were concordant with GeT-RM results for all samples and genes. This indicates that the long-read capture data can be used to accurately assign star (*)-alleles according to current clinical guidelines. However, it is important to note that GeT-RM star(*)-alleles are only based on a limited set of variants that are used in their variant to star (*)-allele translations. To use Aldy 4 for the assignment of CNV’s (Copy Number Variants), a CN neutral region is required, and this should be taken into account when designing a sequencing panel as proposed here. In the absence of a CN neutral region, CNVs for genes like *CYP2D6* cannot be accurately assigned. As the data can still be noisy at lower coverage regions in the *CYP2D6* region it can be expected that the calling of structural variants is not yet optimal, which we have also observed since the star (*)-alleles discrepancies were mainly found in CYP2D6 structural variants.

In conclusion, we have presented a PacBio HiFi long-read sequencing-based approach for pharmacogenomics resulting in high on target phasing and high accuracy calls for both SNVs and clinical star(*)-alleles.

